# Comprehensive characterization of single cell full-length isoforms in human and mouse with long-read sequencing

**DOI:** 10.1101/2020.08.10.243543

**Authors:** Luyi Tian, Jafar S. Jabbari, Rachel Thijssen, Quentin Gouil, Shanika L. Amarasinghe, Hasaru Kariyawasam, Shian Su, Xueyi Dong, Charity W. Law, Alexis Lucattini, Jin D. Chung, Timur Naim, Audrey Chan, Chi Hai Ly, Gordon S. Lynch, James G. Ryall, Casey J.A. Anttila, Hongke Peng, Mary Ann Anderson, Andrew W. Roberts, David C.S. Huang, Michael B. Clark, Matthew E. Ritchie

## Abstract

Alternative splicing shapes the phenotype of cells in development and disease. Long-read RNA-sequencing recovers full-length transcripts but has limited throughput at the single-cell level. Here we developed single-cell full-length transcript sequencing by sampling (*FLT-seq*), together with the computational pipeline *FLAMES* to overcome these issues and perform isoform discovery and quantification, splicing analysis and mutation detection in single cells. With *FLT-seq* and *FLAMES*, we performed the first comprehensive characterization of the full-length isoform landscape in single cells of different types and species and identified thousands of unannotated isoforms. We found conserved functional modules that were enriched for alternative transcript usage in different cell populations, including ribosome biogenesis and mRNA splicing. Analysis at the transcript-level allowed data integration with scATAC-seq on individual promoters, improved correlation with protein expression data and linked mutations known to confer drug resistance to transcriptome heterogeneity. Our methods reveal previously unseen isoform complexity and provide a better framework for multi-omics data integration.

## Main

Single-cell RNA-sequencing (scRNA-seq) is a widely adopted method for profiling transcriptomic heterogeneity in health and disease^1^. However, assessing transcript-level changes between cell types using current scRNA-seq protocols is challenging due to their reliance on short-read sequencing. Previous studies using plate-based methods^2,3^ have focused on individual alternative splicing events such as exon skipping, due to the fundamental limitation of short-read sequencing in linking distal splicing outcomes belonging to the same transcript. The Smartseq3 protocol^4^ can achieve full-length transcript coverage but is still unable to assemble the complete transcript sequence and is heavily reliant on the reference annotation. Droplet-based methods^5^ such as 10x only sequence the 3’ or 5’ end of transcripts which largely precludes isoform identification. Long-read sequencing can overcome this limitation and generate full-length transcript information in single cells, as illustrated in several recent studies^6–9^. However, the throughput of current long-read sequencing platforms is still not comparable to short-read platforms and the per-base accuracy is also lower, which together create many issues. Limited sequencing throughput introduces a trade-off between the per-cell sequencing depth and the number of cells or genes processed. Protocols such as ScISOr-Seq^6^ perform shallow sequencing per cell while RAGE-Seq^7^ focuses on specific transcripts rather than the whole transcriptome. On top of the current protocol limitations, another pressing issue is the lack of data analysis pipelines for long-read transcriptome data, especially for single cells. Methods such as *FLAIR*^10^ and *TALON*^11^ have been developed for isoform annotation and quantification but have not been benchmarked and lack the ability to perform quantification at the single-cell level. Therefore, we need both new protocols and computational pipelines to overcome these limitations.

To this end, we developed FLT-seq and *FLAMES* to perform single-cell isoform sequencing and data analysis. Adapted from the popular 10x Chromium platform, FLT-seq is a cost-effective approach to discover and quantify isoforms in single cells by integrating data from short- and long-read sequencing technologies. By subsampling single cells from a full 10x run and applying nanopore long-read sequencing, FLT-seq can achieve comparable sequencing depth per cell to that obtained from short-read platforms. For data analysis, we developed a computational framework to perform single-cell full-length analysis of mutations and splicing (*FLAMES*), which includes cell barcode, UMI assignment from nanopore reads and semi-supervised isoform discovery and quantification. We applied FLT-seq and *FLAMES* to human and mouse samples containing different cell types and highlight shared splicing patterns in human cancer cells and mouse quiescent muscle stem cells. Differential transcript usage analysis pinpointed common functional modules and genes across samples. We also demonstrate that FLT-seq is a promising tool for detecting coding variants of clinical relevance. Taken together, our new protocol and data analysis pipeline enable comprehensive characterization of the full-length isoforms present in single cells that are currently missed by short-read sequencing datasets.

### High-throughput single-cell full-length transcriptome sequencing with FLT-seq

FLT-seq is based on the Chromium scRNA-seq platform (10x Genomics), with optimizations to better amplify the full-length cDNA. Since the throughput of long-read platforms is still limited compared to Illumina sequencing platforms, we developed a strategy to subsample 10-20% of the 10x Chromium generated Gel Bead-in-Emulsions (GEMs) after reverse transcription (Figure 1A). This is equivalent to sampling 10-20% of the cells as the cDNA of each cell is still within the GEMs. After subsampling, the GEMs are pooled separately for library preparation. Part of the amplified cDNA from the 10-20% subsample is used for Oxford Nanopore Technologies long-read library preparation and sequencing on a PromethION. The remainder of the cDNA from the 10-20% sample together with the GEMs from the 80-90% sample are used for regular 10x library preparation and Illumina sequencing in parallel. In the end, long-read data from the 10-20% subsample of cells and Illumina short-read data for all cells are generated by this protocol.

**Figure 1.**
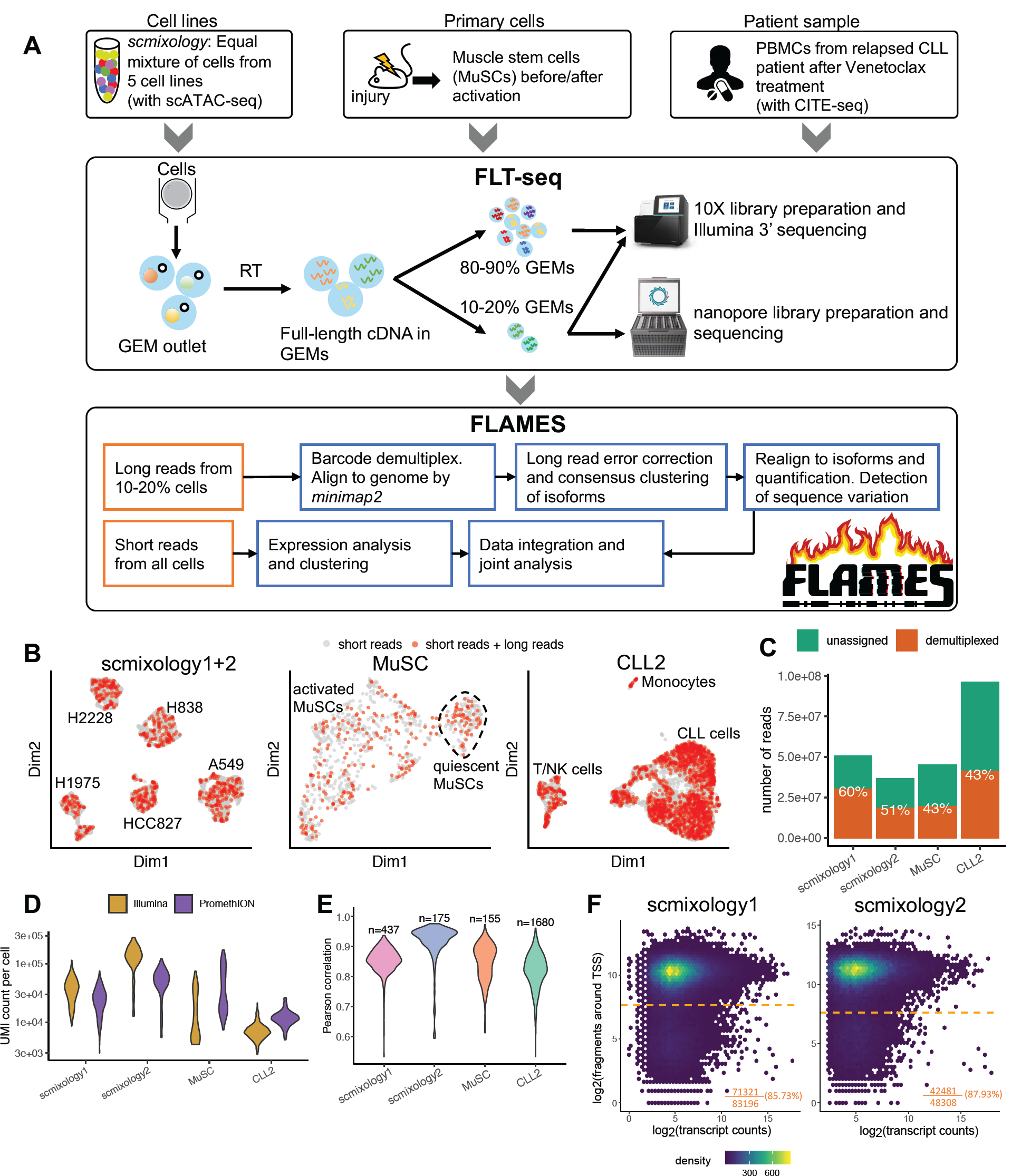
Overview of experimental design, FLT-seq and *FLAMES* methods and basic summary statistics. (A) Summary of the study design, with an overview of the FLT-seq protocol and *FLAMES* data processing pipeline. (B) UMAP visualization of cells in each sample, cells colored in red are sampled for long-read sequencing. *scmixology1* and *scmixology2* were integrated and shown together in one plot. All UMAP visualizations are based on short-read data (C) The number of nanopore reads generated from each sample, and the percentage of reads that were assigned a cell barcode. (D) Distribution of UMI counts per cell for Illumina and nanopore data in each sample. (E) Correlations between gene UMI counts generated from nanopore long-read and Illumina short-read data. (F) Density scatter plot shows the relationship between transcript-level counts and scATAC-seq read counts around the TSS regions. The horizontal orange line shows the threshold calculated that separates open chromatin from the background. The percentage shows the transcripts that have their TSSs in open chromatin regions.

We demonstrate FLT-seq by profiling 16,660 cells from diverse biological systems, 2,737 of which were sequenced by both long-read and short-read technologies (Figure 1A, Table S1). Firstly, we used our previously published *scmixology* design^12^, which involved an equal mixture of cells from five cell lines (H2228, H838, H1975, HCC827, A549). Two biological replicates were profiled (*scmixology1* and *scmixology2*) with FLT-seq together with 10x scATAC-seq for the second replicate. In addition to the cell-line mixtures, we sequenced freshly isolated quiescent and activated muscle stem cells (*MuSC*s) from mouse. Lastly, we applied FLT-seq to a cryogenically preserved peripheral blood mononuclear cell (PBMC) sample from a patient (*CLL2*) whose chronic lymphocytic leukemia (CLL) had progressed on venetoclax treatment after a durable response. The sample was prepared together with the 10x CITE-seq assay with 17 antibody markers. This demonstrated the broad utility of FLT-seq, which is compatible with different 10x transcriptomic assays and can be applied to both fresh and frozen samples. The Uniform Manifold Approximation and Projection (UMAP) visualization presented the cell populations as expected (Figure S1) and revealed no obvious bias in the sampling process of FLT-seq (Figure 1B). Collectively, we generated 230 million long reads across all samples, together with scATAC-seq and CITE-seq data for the *scmixology2* and *CLL2* samples respectively.

### Single-cell isoform detection and quantification with *FLAMES*

We developed a flexible computational framework called *FLAMES* (*<u>F</u>*ull-*<u>L</u>*ength *<u>A</u>*nalysis of *<u>M</u>*utations and *<u>S</u>*plicing in long-read data) to detect and quantify isoforms for both single-cell and bulk long-read data (Figure 1A and Methods). Input to *FLAMES* are fastq files generated from the long-read platform. Using the cell barcode annotation obtained from short-read data as the reference, it identifies and trims cell barcodes/UMI sequences from the long reads. After barcode assignment, all reads were aligned to the relevant genome to obtain a draft read alignment. The draft alignment is then polished and grouped to generate a consensus transcript assembly. All reads are aligned again using the transcript assembly as the reference and quantified.

Next, we benchmarked *FLAMES* on a bulk SIRV spike-in dataset^13^ for which the isoform structure and abundances are known *a priori*. We compared our new method to *FLAIR, TALON* and *StringTie2*^14^ which are designed for bulk long-read isoform detection and quantification. Our results clearly show that *FLAMES* outperforms other tools both in terms of the isoform detection (Figure S2A) and quantification (Figure S2B). It is notable that *TALON* and *FLAIR* still have many false positive transcripts after filtering by abundance (82% and 78% respectively) compared to FLAMES (3%), highlighting the importance of choosing an appropriate method.

After validating the method, we used *FLAMES* to preprocess and analyze the four datasets generated. 40-60% of the long reads could be assigned to an expected cell barcode and were kept for further analysis (Figure 1C). The transcript coverage of reads realigned to assembled transcripts showed that the percentages of full-length reads decreased for longer transcripts (Figure S3), which also been shown in another study^15^. Reads that are not full-length and cannot be uniquely assigned to transcripts are discarded during data processing. The average UMI count per cell ranges from 10,000-60,000, varied by cell type, and was comparable to the short-read counts from the same cells (Figure 1D). Gene-level UMI counts between the matched nanopore and Illumina data were also found to be highly correlated (Figure 1E). The data processed by *FLAMES* showed that FLT-seq generated high quality long-read data that was comparable to the short-read Illumina data when analyzed at the gene-level.

To validate the transcription start sites (TSSs) from the isoforms generated by *FLAMES*, we compared them to the FANTOM5 TSS peaks^16^ and found that around 75% of the TSSs are within the FANTOM5 annotation (Figure S4A). For the *scmixology* data where scATAC-seq data from the same populations were also available, we aggregated scATAC-seq signals around the TSSs as an indicator of open promoters. The result showed *scmixology1* and *scmixology2* have similar open promoters and more than 85% of the TSSs are within active promoters, supporting the existence of these transcripts (Figure 1F). In contrast, when we process the *scmixology* data using *TALON, FLAIR* and *StringTie2* the results were less optimal. The majority of transcripts generated by *FLAIR* and *TALON* did not match the reference annotations (Figure S4B) and *FLAIR, TALON* and *StringTie2* had fewer TSSs overlapping the FANTOM5 annotations compared to *FLAMES* (Figure S4C). We found similar results when comparing the scATAC-seq signals around the TSS regions from transcripts generated by different methods (Figure S4D), with *TALON* and *FLAIR* having around 40% and 50% of their TSSs in open chromatin regions respectively. In summary, our comparisons yield similar results between a spike-in dataset and the *scmixology* dataset, with *FLAMES* outperforming *StringTie2, FLAIR* and *TALON* with the latter two generating many false transcripts.

### Characterization of isoforms reveals the distinct splicing landscapes of different cell populations

We compared the transcripts generated by *FLAMES* to the reference annotation and classified them using the scheme introduced in *SQANTI*^17^, including transcripts with all splice junctions matching to reference transcripts (full splice match, FSM) or partially matching to consecutive splice junctions for a reference transcript (incomplete splice match, ISM) and transcripts with novel splice junctions with new (Novel, not in catalog, NNC) or known (Novel, in catalog, NIC) donor and acceptor sites (Figure 2A). We observed that the number of transcripts detected varies between samples and was correlated with the sequencing throughput as shown in Figure 1C. While around half of the transcripts detected were novel, the majority of reads aligned to known transcripts with novel transcripts having lower abundance in general. In addition to the comparison with reference annotations, we also compared the transcripts generated from the three human samples to each other (Figure 2C) and found many transcripts unique to each sample (22%, 27% and 68% for *scmixology1, scmixology2* and *CLL2* respectively). A majority of transcripts (∼60%) were shared between biological replicates and among these 30% were novel, suggesting that many conserved alternative splicing events in these cell lines have not been annotated.

**Figure 2.**
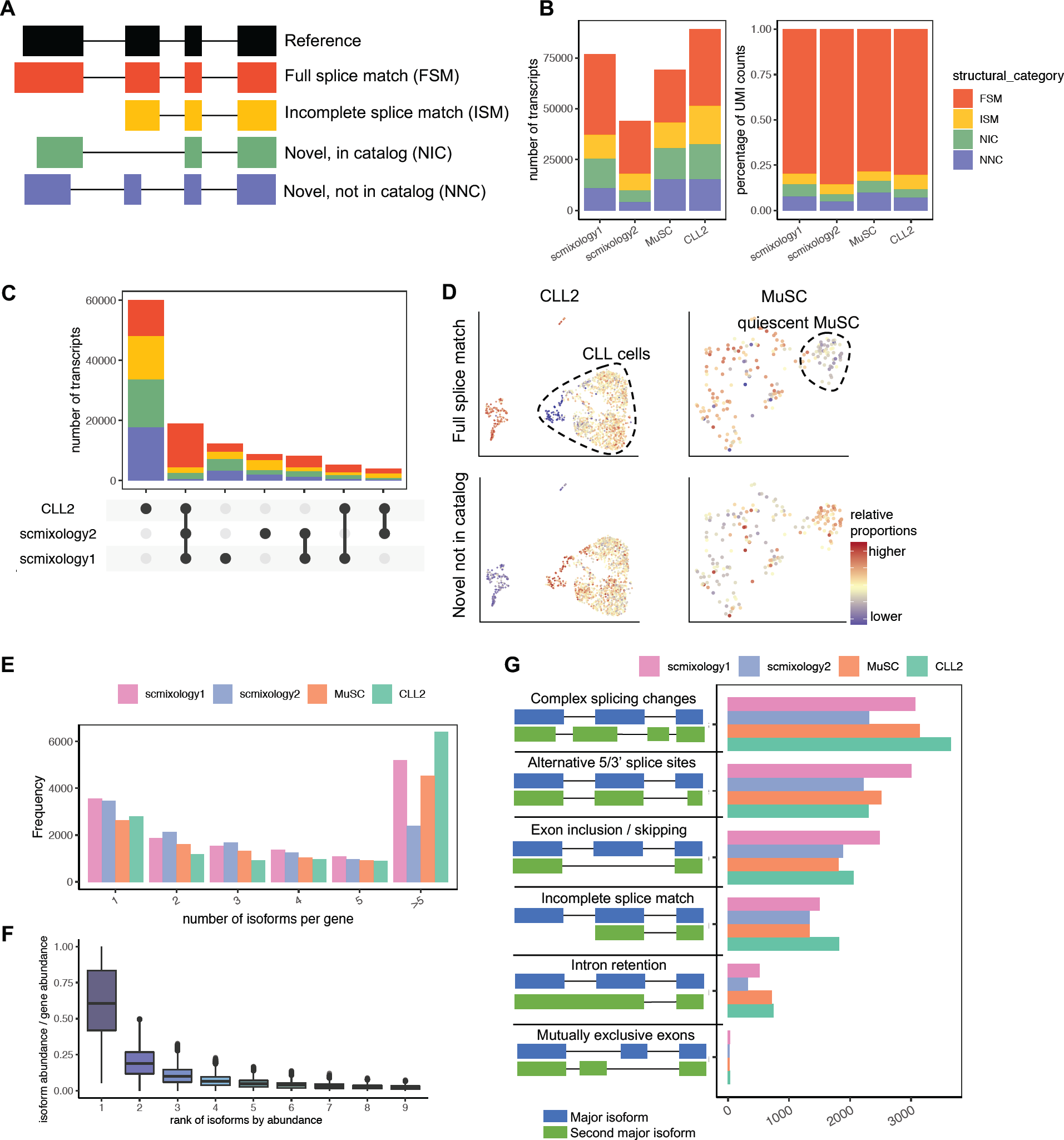
Overview of the single-cell isoform-level analysis from *FLAMES.* (A) Classification of transcripts according to their splice sites when compared to reference annotations. (B) Summary of transcripts in different categories in (A) both in numbers (left) and in the percentage of UMI counts (right). (C) UpSet plot showing overlap of transcripts in human datasets, where number of transcripts shared by different sets of samples are indicated in the top bar chart, colored by categories specified in (A). (D) UMAP visualization of *CLL2* and *MuSC* dataset on the cells sampled for long-read sequencing. Colored by percentage of UMI counts of transcripts in FSM (top) and NNC (bottom) categories. CLL cells and quiescent MuSCs are annotated on the plot. (E) Bar plot of number of distinct transcripts expressed per gene. Genes with more than five distinct transcripts are merged. (F) Box plot showing the percentage of transcript abundance relative to gene abundance for genes express multiple transcripts. Transcripts are ranked by abundance, shown on the x-axis. (G) Summary of the type of alternative splicing between the two most abundant transcripts of each gene. The ‘Complex splicing changes’ category represents transcripts with more than one type of constitutive alternatively spliced event.

Within samples we found consistent alterations in splicing patterns between cell populations (Figure 2D). CLL cells had higher proportions of novel transcripts, especially transcripts with novel splice junctions (NNC), compared to non-CLL cells from the same sample, including T cells, NK cells and monocytes. Similarly, quiescent muscle stem cells also had higher proportions of novel transcripts compared to activated stem cells. Analysis of intronic reads from RNA-seq data^18^ has demonstrated that intron retention, which would produce novel transcripts, is increased in quiescent mouse muscle stem cells and is essential for these cells to exit the quiescent state. This is consistent with our results and suggests that differences in splicing patterns between different cell populations may act as a regulatory mechanism.

After comparing the transcripts identified against reference annotations, we sought to characterize isoforms within the same gene. Around 80% of genes can be expressed as multiple isoforms. The average number of isoforms expressed per gene ranges between 3 and 6 and varied between the different samples (Figure 2E). The distribution of isoform expression is skewed with only a few abundant transcripts dominantly expressed for most genes (Figure 2F). On average, the two most highly expressed isoforms account for 80% of the total counts (median 85%). Next, we categorized the types of alternative splicing between the two most highly expressed isoforms (Figure 2G). Alternative splicing has mostly been studied based on particular events such as exon skipping or alternative 5’ splice sites using short-read sequencing technology^19^. Here we found that around 30% of genes have more than one type of alternative splicing event between the top two isoforms. This means that the 2 most highly expressed isoforms differ by complex splicing changes involving multiple exons, which may not be accurately characterised by short reads because two isoforms could have a skipped exon near the 5’ end and a different splice site near the 3’ end.

In addition to summarizing isoforms at the sample-level, we examined how splicing variants are captured at the single-cell level. Individual alternative splicing events in single cells might follow a unimodal binomial distribution and a cassette exon is skipped or included at the same probability for all cells. Or cells may not share the same parameter of the binomial distribution and create the bimodal splicing pattern, with some cell always skipping an exon while others always include it. The latter is supported by an early single cell study^2^. Recently by analyzing the splice junctions using short-read scRNA-seq data, splicing in homogeneous cell populations has been found to be largely unimodal^20^. Here we expanded the binomial model from considering single splice junctions to complete isoforms in order to estimate the isoform choice between the major isoform and other isoforms (see Methods). These results showed the percentage of cells with binary isoform choice is strongly related to the gene abundance and can be approximated by a unimodal binomial distribution (null distribution), which indicates the cell randomly chooses isoforms with a certain frequency (Figure S5A). We also noticed that the *MuSC* and *CLL2* samples contained more genes with binary splicing patterns, especially for genes with low abundance. This could result from the difference in mRNA amounts where the cell lines are transcriptionally active, and the other samples have lower numbers of mRNA molecules per cells. Or it could be that the *MuSC* and *CLL2* samples have heterogeneous populations with different splicing patterns. To further explore the effect of recovery on isoform identification and splicing analysis, we sampled reads from the *scmixology* dataset to retain 20-80% of the total reads for analysis. The number of transcripts detected decreased with lower sampling rate (Figure S5B), especially for novel transcripts, while binary splicing still showed a unimodal distribution (Figure S5C). This suggested that the reliability of isoform choice estimation may be influenced by multiple factors, including the transcriptional activity and heterogeneity of the cell identities.

### Common classes of genes with differential transcript usage across samples

Following the analysis of isoform abundance, we investigated whether genes with multiple isoforms exhibited differential transcript usage (DTU) between the clusters or cell types shown in Figure S1. We focused on changes in the use of internal splice junctions, grouping transcripts with the same intron chain. To mitigate the high dropout rate in single-cell data, we aggregated the transcript counts into pseudo-bulk values per cluster and filtered out transcripts with low abundance. Next, we evaluated how sequencing level influenced the results of a DTU analysis using the same downsampling scheme used in the last section (see Methods). We observed that although as expected the number of genes with DTU decreases with decreasing sequencing depth per cell, 58% of the changes could still be detected with 20% of the reads (Figure S5). We then performed DTU analysis on all samples and identified between 500 and 1,000 genes with differential transcript usage (gDTU) in each sample (Figure 3A, Table S2). Nearly half of the gDTU (244 out of 573) are shared between *scmixology1* and *scmixology2*. Besides this overlap, gDTU were largely unique for each sample (Figure S6A). The functional annotation clustering of these genes revealed shared pathways across different samples, especially pathways related to transcription and translation, such as mRNA splicing and ribosome biogenesis (Figure 3B, Table S3). As an example, different gDTU related to mRNA splicing were found in different samples. PQBP1, which is involved in pre-mRNA splicing, is a gene with DTU in the *scmixology* samples (Figure S6B), while SRSF2 and SRSF3, which belong to a family that act both as general splicing factors and as regulators of alternative splicing^21^, exhibit DTU in the *CLL2* sample (Figure S6C).

**Figure 3.**
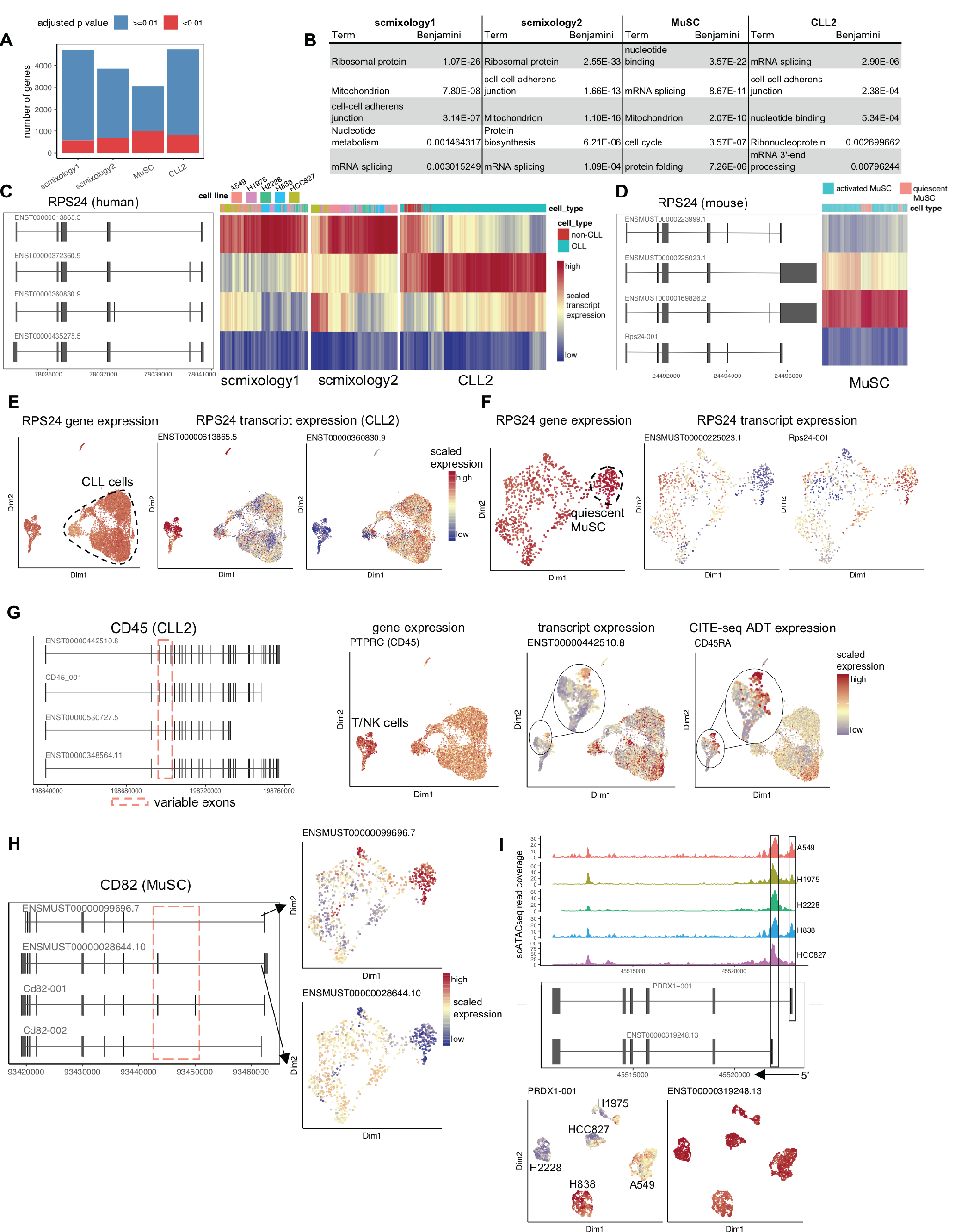
Summary of differential transcript usage results from *FLAMES*. (A) Summary of results from the statistical testing of DTU detected many significant genes per sample (adjusted *P*-value < 0.01). (B) Table of common functional categories among different samples from the functional enrichment analysis of gDTU. (C) Top 4 most abundant isoforms of RPS24 in human and heatmap of their expression at the single-cell level in the *scmixology1, scmixology2* and *CLL2* samples. (D) Top 4 most abundant isoforms of RPS24 in mouse and heatmap of their expression at the single cell level in *MuSCs*. (E) UMAP of cells in *CLL2*, colored by RPS24 gene expression and transcript expression. Two transcripts with differential expression on different populations were selected. Transcript expression in each cell is colored by scaled relative expression to highlight the difference between different populations. (F) Similar to (E), UMAP of cells in *MuSC* sample, colored by RPS24 gene expression and transcript expression. (G) Top 4 most abundant isoforms of CD45 in *CLL2* and UMAP visualizations of the cells colored by (from left to right) gene expression, transcript expression and corresponding protein expression. (H) Top 4 most abundant isoforms of CD82 in *MuSC*, with UMAP visualization of cells colored by expression of two isoforms that have differential expression between quiescent and activated *MuSC.* (I) scATAC-seq read coverage for PRDX1 with cells from each cell line aggregated and plotted together. UMAP plots showing isoform expression, with each cell colored by scaled transcript expressio

Apart from the gDTU that are unique to each sample, we also found a few genes (18) that appeared in all samples (Figure S6A), including a ribosomal protein RPS24 that had the smallest *P*-value among different samples (Table S2). RPS24 is a highly conserved gene between mouse and human, with exon 5 and 6 alternatively spliced to generate different protein coding sequence (Figure 3C, 3D). Further analysis showed transcript usage of this gene is altered at multiple levels in different samples. Firstly, the major transcript of RPS24 was distinct between different samples. Transcripts without exon 5 and 6 were most abundant in the cell lines (*scmixology1* and *scmixology2*, Figure 3C), while the transcript with only exon 5 was highly expressed in the patient derived PBMC sample (*CLL2*, Figure 3C) and *MuSC* sample (Figure 3D). In addition to the major transcript, additional transcripts showed differential transcript usage in different cells within the same sample. The transcript with exon 5 and 6 was more frequently expressed in CLL cells while the transcript without exon 5 and 6 was preferentially expressed in non-CLL cells (Figure 3E). Besides these known transcripts, we also identified a new isoform in mouse (Rps24-001) that was preferentially expressed in quiescent MuSCs (Figure 3F). Different RPS24 isoforms have tissue-specific expression^22^ and some have been linked to tumor progression^23^, however the functional differences between the encoded proteins remains unclear. In summary, we have highlighted the heterogeneity of expression of the RPS24 transcript that is missed in a typical gene-level analysis (Figure 3E, 3F).

Another category of gDTU of interest is cell-surface proteins. Genes encoding cell-surface proteins produce alternative mRNA isoforms, usually by changing the combinations of consecutive exons corresponding to certain functional domains. Some of the isoforms have been characterized, such as CD45, where alternative splicing of exons 4 to 6 are expressed in different lymphocytes^24^. By analyzing the *CLL2* data where the surface marker expression is available through CITE-seq, we detected multiple isoforms of CD45 (Figure 3G). We found similar expression patterns between the transcript and the protein it encodes, where the protein was quantified by counting antibody derived tags (ADT) from CITE-seq data. The result both validated the isoform quantification from the *FLAMES* pipeline and showed that transcript-level analysis can provide better correlation between mRNA and protein expression that cannot be achieved using gene-level quantification. We also observed other genes encoding important cell surface markers with DTU, such as CD82 (Figure 3H), CD47 (Figure S6D) and CD44 (Figure S6E), with different combinations of consecutive exons. Notably, some of the novel transcripts in CD47 can be observed in different samples (Figure S6D), suggesting conserved alternative splicing patterns that are missing from the reference annotation. In sum, we observed previously overlooked isoform diversity of cell-surface proteins, which could introduce functional diversity and contribute to development and diseases^25,26^.

Alternative promoters have been shown to regulate cancer-specific transcription^27^. Here we found transcripts with different TSSs expressed in different cancer cell lines. We have shown that open chromatin captured by scATAC-seq is correlated with the promoter region indicated by the TSS (Figure 1F). As an example, we found multiple isoforms of PRDX1, including a new isoform PRDX1-001 with a different TSS. The new isoform expressed in H838 and A549 contains a novel first exon that is not found in the other cell lines (Figure 3I). Additional open promoter regions are observed in the scATAC-seq data from H838 and A549 that coincide with the new exon (Figure 3I). This result suggests that FLT-seq can be integrated with scATAC-seq at the promoter level to further enhance the resolution of integration and reveal promoter heterogeneity that cannot be found via gene-level integration^28^.

### FLT-seq links coding sequence variation to transcriptome heterogeneity

Nanopore long-read sequencing allows full transcript coverage compared to 3’ or 5’-end short-read sequencing, which provides us with a better chance of identifying coding variations. But the high error rate in sequencing presents an obstacle^29^. To overcome this challenge, we first exclude homopolymer regions since they have higher sequence error rates^30,31^. After excluding these regions, ∼95% of regions have a reference allele frequency >90% (Figure S7A). The remaining single-nucleotide variants (SNVs) were filtered again using a statistical test in *FLAMES* to enrich for true positives and clonal mutations, based on the assumption that the occurrence of sequencing errors is independent of cell barcode and will occur randomly in all cells (Figure 4A and Methods). We tested our approach on the *scmixology* samples where the five cell lines carry distinct mutations. The principal-component analysis (PCA) on the filtered allele count matrix successfully recapitulated the expected population structure (Figure 4B). Louvain clustering on leading principal components generated similar results to the cell type assignment obtained from running *Demuxlet*^32^ on the short-read data (98.7% and 99.4% concordance for *scmixology1* and *scmixology2* respectively), which indicated that we can successfully capture mutations in these different cell lines using long-read data. We then performed differential allele frequency analysis to find SNVs that were specific to each cell line. The results showed a high precision (80.1%) with the SNVs called from bulk RNA-seq data from individual cell lines.

**Figure 4.**
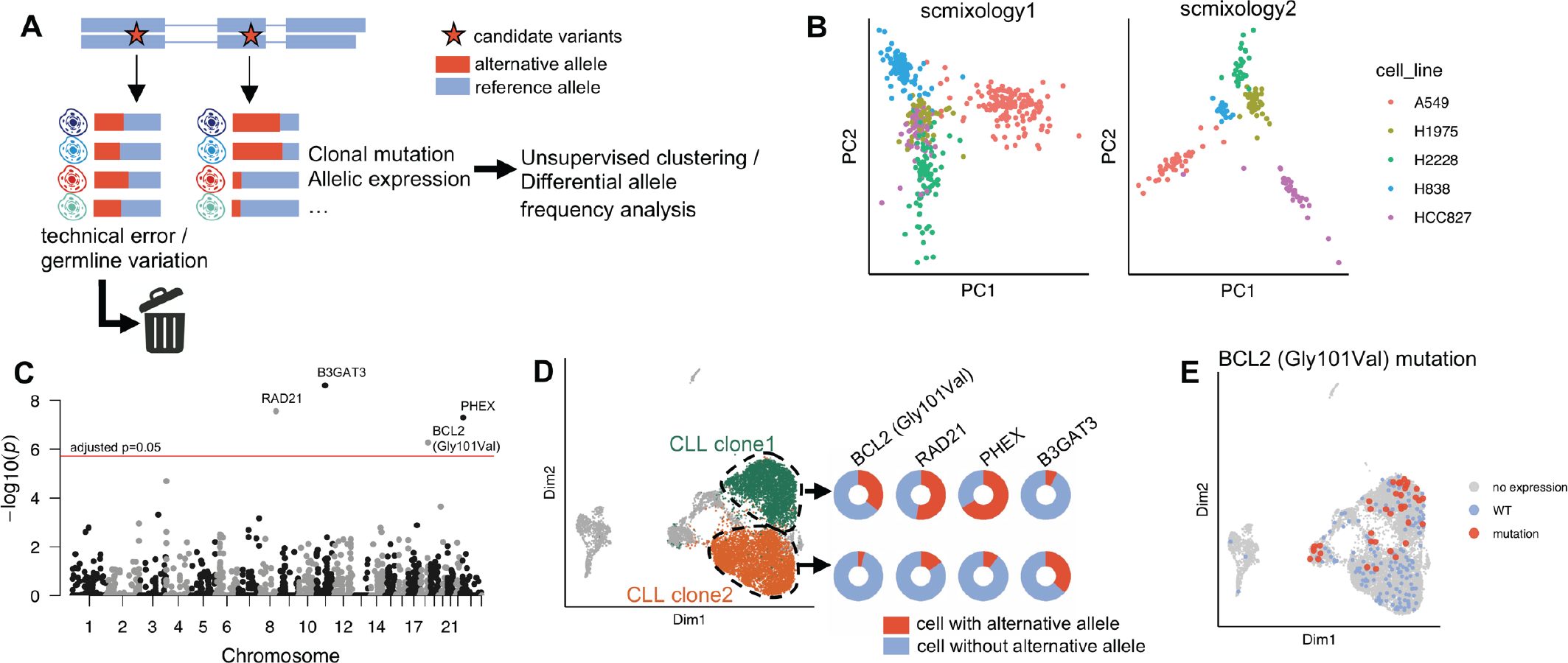
Summary of differential allele frequency analysis to detect coding mutations. (A) Summary of variation filtering and analysis pipeline implemented in *FLAMES*. Candidate variants are filtered based on allele frequency first, then based on per cell allele frequency to remove technical artefacts. The remaining variants are used for differential allele frequency analysis. (B) PCA on alternative allele matrix using the variants after filtering, colored by unsupervised clustering results using top PCs and annotated with cell lines. (C) Manhattan plot of *P*-values from a differential allele frequency analysis with Benjamini–Hochberg adjustment. Genes with significant variants are labelled. (D) UMAP visualization highlighting two CLL populations that have differential allele frequency for the significant variants. (E) UMAP visualization of cells colored by Gly101Val mutation status.

After we confirmed the approach using the cell lines, a similar analysis was performed on the *CLL2* dataset to examine the relationship between transcriptome heterogeneity and sequence variations. We searched for SNVs that only existed in the cancer cells and had a differential allele frequency across different CLL transcriptional clusters. We found four significant mutations (adjusted *P*-value < 0.05, chi-square test, Benjamini– Hochberg correction) associated with different CLL clusters (Figure 4C, 4D) and their existence was confirmed by bulk genomic sequencing (data not shown). By investigating the allele frequency of these mutations across clusters (Figure S7B), we identified two subclones, where mutations in the genes BCL2, RAD21 and PHEX are enriched in *subclone1* and the mutation located in the gene B3GAT3 is enriched in *subclone2* (Figure 4D). Although the analysis itself does not require genomic sequence data as a reference, it is important to validate the mutations detected since the changes in allele frequency may also come from allele-specific expression or RNA editing. The Gly101Val mutation has been confirmed to promote resistance to venetoclax treatment by reducing the affinity of BCL2 for venetoclax^33^, and patient *CLL2* was known to carry ∼25% Gly101Val mutations. Using FLT-seq, we showed that the mutation is not just subclonal, but also linked to specific transcriptional clusters. Taken together, FLT-seq provides unbiased high-throughput linking of single-cell mutations and transcriptomic heterogeneity.

## Discussion

Transcriptional activity is typically summarized at the gene-level due to the limitations of short-read technology, especially in scRNA-seq studies. The recent development of long-read sequencing technology promises sequencing of the full-length transcript, but its application to single cells has been slowed by a lack of protocols and data analysis pipelines. By optimizing the 10x Genomics scRNA-seq protocols we have developed the scalable and affordable FLT-seq approach by cell sampling to achieve per-cell sequencing depths similar to short reads at a much lower cost than other methods^6^. It combines the advantages of throughput and accuracy from short-read sequencing to identify cell barcodes and perform clustering with the strengths of long-read sequencing that enables isoform discovery and quantification, the results of which are combined in an integrated analysis. FLT-seq is compatible with a variety of 10x transcriptomic assays and could be potentially applied to any single-cell transcriptomic protocol with cell barcoding, including Visium spatial transcriptomics. It could also be combined with other single-cell long-read sequencing approaches such as R2C2^34^ or other sequencing platforms including PacBio Sequel II. As the throughput and accuracy of long-read sequencing platforms increases, it may be possible to skip FLT-seq’s sampling step and apply long-read sequencing only rather than the current hybrid approach to achieve similar aims in future studies.

To analyze noisy long-read data, we developed the *FLAMES* pipeline that can detect and quantify novel and known isoforms in single-cell and bulk samples. It can also find cell-type-specific isoforms and mutations through comparative analysis. We showed that *FLAMES* outperforms other methods both on a synthetic spike-in dataset^35^ and experimental datasets. Our comparison indicated that certain methods generate many spurious transcripts, highlighting the importance of designing appropriate methods to detect novel isoforms.

With FLT-seq and *FLAMES*, we performed the first comprehensive characterization of full-length isoforms at the single-cell level across species and cell types. We detected thousands of novel transcripts expressed at low levels, most of which were unique to each sample. Novel transcripts also showed distinct enrichment patterns at the single-cell level where CLL cells and quiescent muscle stem cells often have more novel transcripts. We frequently found more than one differentially spliced junction between isoforms for many genes, suggesting the inadequacy of studying individual splicing junctions via short reads. Furthermore, we found that single cells often express more than one mRNA isoform per gene and the isoform choice is largely random, challenging the argument that individual cells have binary outcomes in splicing^36^. In addition to the overall stochasticity of isoform choice, we observed cell type specific isoform usage for genes that are enriched for common functions such as mRNA splicing and ribosome biogenesis. The expression of ribosomal and spliceosome proteins has low correlation with their gene-level expression^37^, indicating some protein expression levels might be regulated by alternative splicing. Genes encoding cell-surface proteins such as CD44 and CD47 often have cell-type-specific splice variants, some of which have not been previously annotated, and may result in different functions^36^. The examples we show indicate that identification of cell-type-specific isoforms may have broad applications in many fields such as immunotherapy, where cell surface proteins play an important role.

Compared to short-read scRNA-seq, FLT-seq provides better linkage between proteome and genome as it can resolve the full-length transcript. Transcript abundance can be aligned directly to single-cell protein measurements, which is particularly helpful for proteins with multiple isoforms such as CD45. We also showed that FLT-seq facilitates integration with scATAC-seq data, such that open chromatin signals are not summarized at the gene-level but at the exact transcription start site. Full transcript coverage unlocks the potential to detect mutations and allelic expression for specific isoforms. Through unsupervised analysis, we not only identified coding mutations that cannot be easily detected by 3’ or 5’-end short-read sequencing, but we could also associate mutations to different transcriptional signatures. This can improve our understanding of the correlation between the single-cell transcriptome and genome in cancer.

FLT-seq and *FLAMES* provide new avenues for characterizing single-cell transcriptomic heterogeneity at the transcript-level and unveils new questions and challenges. Benchmarking studies that compare the performance of emerging long-read sequencing protocols and computational methods are needed to help our understanding of the strengths and weaknesses of different approaches. Many unanswered questions about transcript expression and alternative splicing in single cells also remain. For example, do the majority of novel transcripts simply reflect stochastic noise in the splicing machinery^38^, or are they indicative of a genuine increase in protein diversity? Our methods and analysis provide a starting point for addressing these questions to accelerate isoform-level studies in single cells.

## Acknowledgements

We thank Clare Weeden and Marie-Liesse Asselin-Labat for providing the cell lines used in this study, Stephen Wilcox for Illumina sequencing, Kathleen Zeglinski for designing the *FLAMES* logo and Joan Heath for providing critical feedback on this manuscript. This work was supported by funding from the Chan Zuckerberg Initiative DAF, an advised fund of Silicon Valley Community Foundation (grant numbers 2018-182819 and 2019-002443 to M.E.R.), a Melbourne Research Scholarship to L.T, a Leukemia & Lymphoma Society (LLS) of America Fellowship to R.T. (5467-18) and LLS Specialized Center of Research [SCOR] grant no. 7015-18 to A.W.R. and D.C.S.H., a Cure Cancer and Cancer Australia grant (1186003 to R.T.), fellowships and grants from the Australian National Health and Medical Research Council (NHMRC: Project Grant 1143163 to M.E.R.; Program Grants 1016701 to D.C.S.H. and 1113577 to A.W.R.; Research Fellowships 1079560 to A.W.R., 1156024 to D.C.S.H. and 1104924 to M.E.R.; Investigator Grant 1174902 to A.W.R.), the Australian Research Council (Discovery Project 200102903 to M.E.R.), the Genomics Innovation Hub, Victorian State Government Operational Infrastructure Support and Australian Government NHMRC IRIISS.

## Author contributions

L.T. developed the *FLAMES* package, performed experiments, conducted data analysis and wrote the manuscript. J.S.J. developed the FLT-seq protocol and performed experiments. R.T., M.A.A, H.P. and C.J.A.A collected patient samples and performed experiments related to *CLL2*. C.H.L., J.D.C., T.N. and A.C. performed experiments related to *MuSC*. Q.G., H.P. and A.L performed nanopore sequencing. S.L.A., H.K., X.D. and C.L. analyzed data and S.S. developed software. G.S.L, J.G.R., A.W.R. and D.C.S.H. facilitated the *MuSC* and *CLL2* studies and provided helpful comments and discussion. J.S.J, Q.G and M.B.C. planned the research and provided helpful comments and discussion. M.E.R. planned and supervised the research and wrote the manuscript. All authors read and approved the final manuscript.

## Methods

### Human Cell Lines

The cell culture and sample preparation of the *scmixology* cell lines are as previously described. Briefly the five cell lines (H2228, H838, H1975, HCC837, A549) were retrieved from the ATCC (https://www.atcc.org/) and cultured in Roswell Park Memorial Institute (RPMI) 1640 medium with 10% fetal calf serum and 1% penicillin-streptomycin. The cells were grown independently at 37°C with 5% carbon dioxide until near 100% confluency. The cells were then counted and mixed in equal numbers. The mix was used for FLT-seq library preparation. The first batch was derived from the same sample featured in a previous study (*scmixology1*). The cell lines were cultured again using the same protocol to create a second batch (*scmixology2*) processed by both FLT-seq and scATAC-seq.

### Mouse Muscle Stem Cells

#### Animals

All procedures were approved by the Animal Ethics Committee of The University of Melbourne and conformed to the Australian code of practice for the care and use of animals for scientific purposes as stipulated by the National Health and Medical Research Council of Australia. Mice were housed in the Biological Research Facility at The University of Melbourne under a 12h light-dark cycle, with drinking water and standard chow provided ad libitum. Pax7^creERT2^R26-eYFP^fl/fl^ mice were generated from Pax7^creERT2^ and R26-eYFP^fl/fl^ founder mice strains^39,40^ both on a C57BL6 background.

#### Muscle Injury and Stem Cell Isolation

At 3 months of age, Pax7^creERT2^; R26R^YFP^ mice received daily 100µL tamoxifen (20mg/ml in corn oil) for five days to label MuSCs with YFP. Two weeks after the final injection, animals were killed and hindlimb muscles excised and dissociated as described previously^41^. Cells were sorted on a FACS Aria III (BD Biosciences) with gating based on YFP.

To obtain activated MuSCs, mice were anaesthetized with isoflurane and muscles were injured with 1.2% barium chloride (Sigma). Lower hindlimbs received 40µL barium chloride via an intramuscular injection. Activated cells were isolated 72 hrs post injury while quiescent MuSCs were isolated from uninjured muscles as described previously.

### Human CLL Sample

After providing written informed consent, the patient sample was collected after progression on venetoclax treatment^42^ (Human Research Ethics Committee approvals: Melbourne Health 2011.044, 2016.305; Peter MacCallum Cancer Centre 11/18; Walter and Eliza Hall Institute 05/04). Blood was collected in EDTA tubes and processed within 2 hours. Peripheral blood mononuclear cells (PBMCs) were isolated using Ficoll-Paque Plus (#17144002, lot:10258101, GE Healthcare) density gradient centrifugation and cells were cryopreserved. PBMCs were thawed, rested for 2 hours, and incubated with Fc Receptor blocking solution (Human TruStain FcX, Biolegend) for 10 minutes prior to staining with TotalSeq C antibodies (Biolegend) at 4 degrees for 30 min. PBMCs were washed three times and stained with propidium iodide (PI, Sigma). Viable cells (PI negative) were flow sorted using the FACSAria (BD) and diluted to 1000 cells/µL.

### FLT-seq

Single-cell capture and cDNA amplification for mouse skeletal muscle stem cell and cell lines was performed using 10x Genomics Chromium Single cell 3’ Library and Gel Bead (v2 for *MuSC* and *scmixology1* and v3 for *scmixology2*) and for *CLL* sample using Chromium Single Cell 5′ Library & Gel Bead Kit (v1) according to manufacturer instructions. Full-length cDNA generation for FLT-seq was carried out as described in detail at protocols.io^43^. Briefly we followed the standard 10x genomic user guide, with RT time increased to 2 hours to increase reverse transcription of longer transcripts. After GEM-RT we transferred 10-20% volumes of GEMs into a new tube and performed subsequent steps in parallel for both 10-20% and 80-90% subsample where each subsample is treated as a separate sample according to the 10x user guide. Volume of the Cleanup Mix was reduced accordingly. cDNA generation for hybrid capture input was similar to FLT-seq except that cDNA from the remaining sample (i.e. 80-90%, excluding the 10-20% subsample) was used for amplification using primers 10x Partial R1: CTACACGACGCTCTTCCGATCT and T5’ PCR Primer IIA: AAGCAGTGGTATCAACGCAGAG in place of FPSfilA and RPSfilBr.

Standard single cell Illumina libraries were prepared according to 10x protocol. Illumina HiSeq2500 was used for sequencing *scmixology1* (2×125 cycles) and *MuSC* (2×150 cycles). Other libraries were sequenced on NextSeq 500 (1x 28 / 1×91 cycles plus 8 base index cycle) using the v2 150 cycle High Output kit (Illumina) as per the manufacturer’s instructions. The base calling and quality scoring were determined using Real-Time Analysis on board software v2.4.6, while the FASTQ file generation and de-multiplexing utilised bcl2fastq conversion software v2.15.0.4. Full-length cDNA libraries from subsamples or capture cDNA were prepared using Oxford Nanopore Technologies SQK-LSK109 Ligation Sequencing Kit with the following modifications: incubation times for end-preparation and A-tailing were lengthened to 15 minutes, and all washes were performed with 1.8X Ampure beads to conserve smaller fragments. SFB was used for the final wash of the libraries. 50 fmol per library were sequenced on PromethION FLO-PRO002 R9.4.1 flow cells according to manufacturer protocols.

### Single cell ATAC-seq (scATAC-seq)

The cells from five cell lines were counted and mixed equally. Cell nuclei were isolated and washed according to the Nuclei Isolation for Single Cell ATAC Sequencing (10x Genomics) protocol, with 1 million cells to start with (0.2 million from each cell line) and 3 minutes lysis on 100µL buffer. Nuclei were then used to generate scATAC-seq libraries according to the Chromium Single Cell ATAC Reagent Kits User Guide (10x Genomics; CG000168 Rev B) Sequencing libraries were loaded on an Illumina sequencer with 2 × 75 paired-end kits using the following read length: 72 bp read 1N, 8 bp i7 index, 16 bp i5 index and 72 bp read 2N. In the sequencing reaction, reads 1N and 2N contain the DNA insert, while the index reads, i5 and i7, capture the cell barcodes and sample indices, respectively. Cells were sequenced on Illumina HiSeq2500 with near around 300 million read pairs in total.

### scATAC-seq data analysis

scATAC-seq sequencing data was demultiplexed, preprocessed and aligned with the default settings of the single cell ATAC *Cell Ranger* platform (1.0). The reference used for alignment through the *Cell Ranger* platform was hg38. Next, *Picard tools* (http://broadinstitute.github.io/picard/) *was used to remove the PCR duplicates. Samtools*^44^ (1.7) was used to extract read pairs that have mapping quality (MAPQ)>30, were nonmitochondrial and not chimerically mapped. *bedtools*^45^ (v.2.26.0) was used to identify reads in mate pairs (i.e. fragments) and adjust the start of the paired reads to account for the 9bp region that the transposase enzyme occupies during transposition (i.e. +4 bp for + strand and −5 bp for - strand). Next, *Demuxlet* was used to identify the cell lines of each cell barcode using the genotypic information acquired in our previous benchmarking studies^12,46^.

The count matrix was generated for each barcode separated BAM file using the *featureCounts* function in the *Rsubread* package^47^ (1.32.4) in the *R* environment (v. 3.5.1). The annotation features were promoter regions (i.e. TSS - 500 bp to TSS + 200 bp) corresponding to isoforms identified by the various long-read analysis methods (i.e. *FLAMES, TALON, FLAIR, StringTie2*). To determine the background of scATAC-seq and identify the open promoter, we performed analysis on randomized TSS, where a random position in each gene body was used as the TSS. The 90% percentile of the fragment count around random TSS (7.64) was used as the threshold to determine the open promoters and is annotated in Figures 1F and S4D.

For Figure 3I, scATAC-seq coverage was calculated from the aligned BAM files using *compute_coverage* in the *plyranges* package^48^ (1.7.14) and visualised using *view_coverage* in the *superintronic*49 package (0.99.4).

### Illumina short-read data analysis

The fastq data were processed by *scPipe* to generate a gene count matrix for all samples except *CLL2*, which was processed by *Cell Ranger* (3.0.0) to generate the antibody and gene count matrix. Each gene count matrix was used as input to the standard *Seurat* pipeline with normalization performed by *SCTransform*^50^. Clustering was performed for the *MuSC* and *CLL2* samples with resolution equal to 0.6. The cell line annotation for *scmixology* was acquired using *Demuxlet* with the same parameters as our previous benchmark study. Integration of the *scmixology1* and *scmixology2* datasets was performed using *Seurat*. The clustering results and cell line annotation is shown in Figure S1.

### Nanopore sequencing and data preprocessing

We performed basecalling on the raw fast5 data using *Guppy* (1.8.1 for *MuSC* sample and 3.1.5 for *scmixology* and *CLL2*) from Oxford Nanopore Technologies. For each read, we locate the barcode sequence by searching for the flanking sequence before the cell barcode. The cell barcodes identified from the short-read data provide a reference to search for and trim in the long reads. An edit distance of up to 2 is allowed during cell barcode matching. Reads that failed to match any cell barcode were discarded. Sequences following the cell barcode were used as UMIs and trimmed. For the 3’ end protocol, the polyA tail after the UMI sequence was trimmed and sequences after the polyA tail were kept. The cell barcode and UMI sequence was integrated into the fastq read header as per *scPipe*^51^. The processed fastq was used as input for genome alignment and further analysis.

### Detection and quantification of isoforms

Reads were aligned to the genome by *minimap2*^52^ (-ax splice --junc-bonus 1 -k14 -- secondary=no --junc-bed) using Gencode reference (human hg38.v33, mouse mm10.vM24). *FLAMES* summarizes the alignment for each read by grouping reads with similar splice junctions (<5bp) to get a raw isoform annotation. The raw isoform annotation is compared against the reference annotation to correct potential splice site and transcript start/end errors. Transcripts that have similar splice junctions (<5bp) and transcript start/end (<100bp) to the reference transcript were merged with the reference. This process will also collapse isoforms that are likely to be truncated transcripts. Next, the sequence of each polished transcript was extracted and used as the updated reference. The reads were realigned to this reference by *minimap2*. The transcript coverage of individual reads is summarized in Figure S3. We noticed that the *scmixology2* data contained more reads that were not full-length, which might relate to the difference in sample preparation time or the v2 and v3 10x chemistry. The transcripts with fewer than 5 full-length aligned reads (>95% coverage) were discarded. The reads were assigned to transcripts based on both alignment score, fractions of reads aligned and transcript coverage. Reads that cannot be uniquely assigned to transcripts or had low transcript coverage (<60%) were discarded. The UMI transcript count matrix was generated by collapsing the reads with the same UMI in a similar way to what is done for short-read scRNA-seq data, but allowing for an edit distance of up to 2. Downsampling was performed by randomly selecting reads in bam files that aligned to the genome according to the sampling ratio. The counts of transcripts from the same gene were aggerated to generate the gene-level UMI count and compared to the gene count generated from the short-read data in Figure 1E. *FLAMES* is written in python and uses other packages, including *pysam*^44^ (0.15.2), *numpy*^53^ (1.14.2) and *editdistance* (0.5.3) (pypi.org/project/editdistance).

### Comparison of *FLAMES* to other tools

Direct RNA sequencing data from SIRV spike-in E2 mix which contains 69 synthetic isoform transcripts (from 7 SIRV genes) was downloaded from NCBI (SRX3204589). Alignment was performed using *minimap2* (2.17), with “-ax splice --splice-flank=no - k14 --secondary=no” and “--junc-bed”. For *TALON*, the mapped reads were processed using *TranscriptClean* (1.02) to correct for mismatches and microindels. Following correction, long reads were collapsed into a transcript isoform quantification table in *TALON* (4.1), using the SIRV annotation. We ran the *FLAIR* pipeline (1.4) using default parameters with the SIRV annotation supplied to aid isoform detection. We ran *StringTie2* (2.0.4) with “-L -G -c 10” and uses the SIRV annotation. For *FLAMES*, we used the default parameters except “strand_specific:1” and we filtered transcripts with at least 10 reads. For comparison, transcripts generated by *TALON* and *FLAIR* were also filtered to have a read count of at least 10. The *scmixology1* data was processed in a similar way to the SIRV data. Gencode human hg38.v33 was used as the gene reference annotation for each method. *TALON* was run in parallel on each chromosome to reduce compute time and the results were aggregated later.

### Isoform classification and splicing analysis

*SQANTI2* (4.1, https://github.com/Magdoll/SQANTI2) was used to compare the transcripts identified to the reference with parameter “-g --cage_peak --coverage -- force_id_ignore”. We used the FANTOM5 cage peak dataset on hg19 and mm9 and lifted these to the hg38 and mm10 reference using UCSC’s *liftOver* tool (https://genome.ucsc.edu/cgi-bin/hgLiftOver). The isoform classification was extracted from the *SQANTI2* result and plotted in Figure 2B. The *gffcompare*^54^ (0.11.2) program was used with parameter “-T -R -M” to compare isoform annotations generated from different samples (Figure 2B). It was also used to compare isoform annotations obtained after downsampling (Figure S5B). Results from these comparisons were plotted using *UpsetR*^55^ (1.4.0). We ranked transcript abundance for each gene that had multiple isoforms and obtained the alternative splicing events from the most expressed transcript and the second most expressed transcript. We used a common model to classify the splicing events^56^, where alternative 5/3’ splice site includes alternative promoter and alternative polyadenylation. Transcripts with more than one splicing event were classified as *complex splicing changes* (Figure 2G).

The binary splicing analysis was performed for each multi-isoform gene by calculating the percentage of cells that only expressed the major isoform (isoform with highest abundance) or only expressed other isoforms. The null distribution for each gene assumes each cell randomly chooses isoforms and the chance of picking the major isoform is proportional to the major isoform expression *p*_*m*_, calculated by dividing the major isoform’s total UMI count by the total gene UMI count across all cells. Next, we assume the count of the major isoform follows a binomial distribution *B*(*n, p*_*m*_) where n is the UMI gene count for a cell. Therefore, the probability of binary splicing is Pr(0, *n, p*_*m*_) + Pr (*n, n, p*_*m*_). We used the UMI gene count for each cell Pr(0, *n, p*_*m*_) + Pr (*n, n, p*_*m*_)take the average. In Figure S5A, S5C we fit the relationship between the average binary splicing probability and total gene count (black lines) using *geom_smooth*() in *ggplot2*^56^ (3.3.1) which uses generalised additive models.

### Differential transcript usage analysis

We filtered genes to have at least two isoforms, each with more than 15 UMI counts > 15. For each gene, the per cell transcript counts were merged by group to generate pseudo bulk samples. For scmixology the groups are based on cell line identify inferred by known genetic variation, and for the MuSC and *CLL2* data the groups are based on the clusters acquired from *Seurat* clustering, shown in Figure S1. The top 2 highly expressed transcripts for each group were selected and a UMI count matrix where the rows are selected transcripts and columns are groups was used as input to a chi-square test of independence (*chisq.test* in R). *P*-values were adjusted by Benjamini–Hochberg correction^57^ and results were summarized in Table S2. We performed functional clustering with DAVID^58^ using genes with significant DTU (adjusted *P*-value < 0.01) as input (Table S3). The transcript structures in all figures were plotted using *geom_alignment* in *ggbio*^59^ (1.36.0). We performed imputation of transcript counts on cells that were not sampled for long-read sequencing using the shared nearest neighbour network constructed by *Seurat*^60^ (3.1.5). We then scaled the transcript expression matrix for each gene and the results were used in heatmaps and UMAP visualizations (available in *Seurat)* in Figure 3 and Figure S6. The imputed results were used for visualization purposes only.

### Variant calling and clonal analysis

First, we identified candidate SNVs using *FLAMES* by excluding homopolymer regions (runs >3 of the same nucleotides), positions with coverage of fewer than 100 reads and positions with reference allele frequency less than 10% or greater than 90%. For each candidate SNV, we generated an allele count matrix of the reference and alternative allele. Next, we collect the cells that have reads with the reference allele and the alternative allele and performed a binomial test on the allele counts in a similar way as the binary splicing analysis, assuming that under the null hypothesis each cell has the same probability of having the alternative and reference allele. *P*-values were adjusted by Benjamini–Hochberg correction and candidate SNVs with an adjusted *P*-value < 0.05 were kept for further analysis. PCA was applied to the filtered alternative allele count matrix from the *scmixology* data and first two PCs were plotted in Figure 4B. The top 5 PCs were selected to build the shared neighbour network (*scran::buildSNNGraph*^61^ k=20, d=5) which is used for Louvain clustering (*igraph::cluster_louvain*^62^) in order to examine whether the allele count matrix captured the variants in the cell lines. Next, we performed differential allele frequency analysis in a similar way to the DTU analysis, but with the allele counts aggregated for each cluster. The candidate SNVs with adjusted *P* values < 0.05 were selected and compared to the VCF reference generated from a previous study using bulk RNA-seq. After analyzing the *scmixology* dataset, we processed the *CLL2* data in the same way and conducted differential allele frequency analysis on the *CLL2* clusters shown in Figure S1. The bar plot of cells with alternative alleles of significant SNVs were shown in Figure S7B and summarized in Figure 4D. The capture enrichment data was analyzed in similar way, with the reads processed by *FLAMES* and an allele count matrix generated for the Gly101Val mutation. All plots were generated using *ggplot2* unless otherwise specified and most of the analysis was performed in *R*^63^ (4.0) unless otherwise specified.

## Data and code availability

Raw data are available from GEO under accession numbers GSE126906 and GSE154869 (*scmixology1*), GSE154870 and GSE142285 (*scmixology2*), GSE154868 (*MuSC*) and the patient data (*CLL2*) is available from EGA. Refer to Table S1 for a summary of these datasets. The *FLAMES* source code is available from https://github.com/LuyiTian/FLAMES. The processed data and scripts used in this study are available from https://github.com/LuyiTian/FLTseq_data.

