## Supplementary Figures for "Comprehensive characterization of single cell full-length isoforms in human and mouse with long-read sequencing"

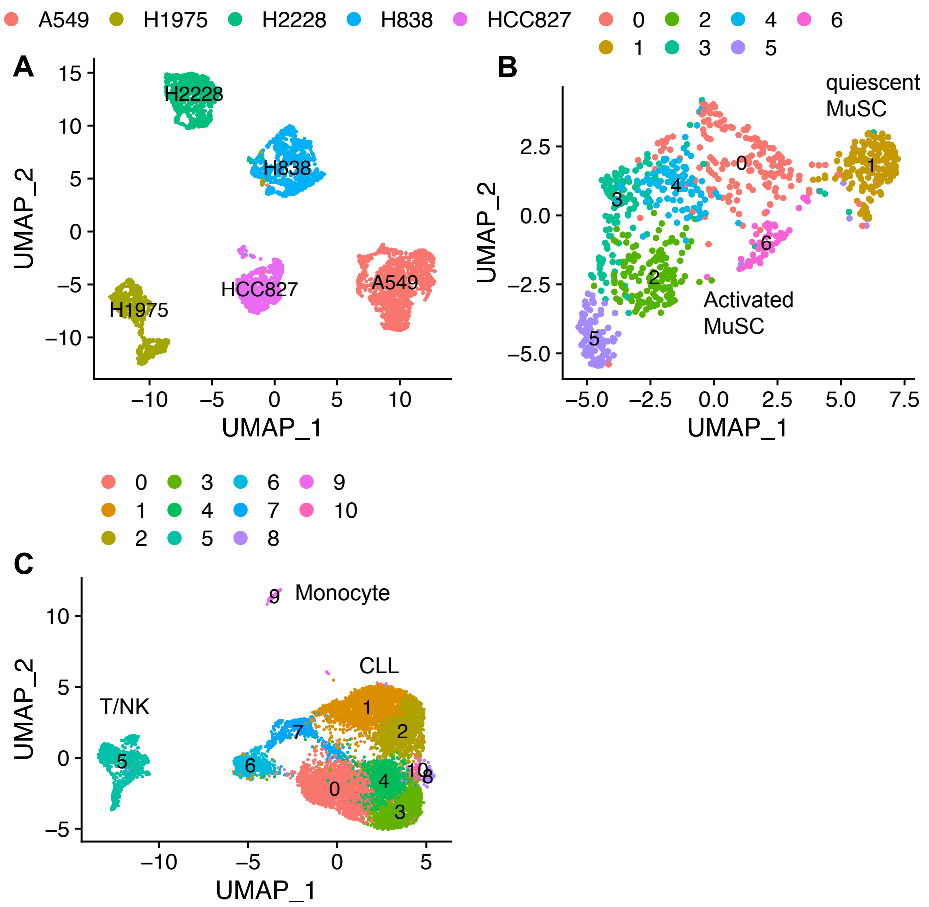


**Figure S1. Dimensionality reduction (UMAP) of all samples.**
UMAP visualization of cells from (A) *scmixology1* and *scmixology2* combined, (B) mouse muscle stem cells (*MuSCs*) and (C) patient *CLL2*. Cells are either colored by cell line identity (*scmixology1* and *scmixology2*) or by *Seurat* clusters and are annotated with major cell populations.

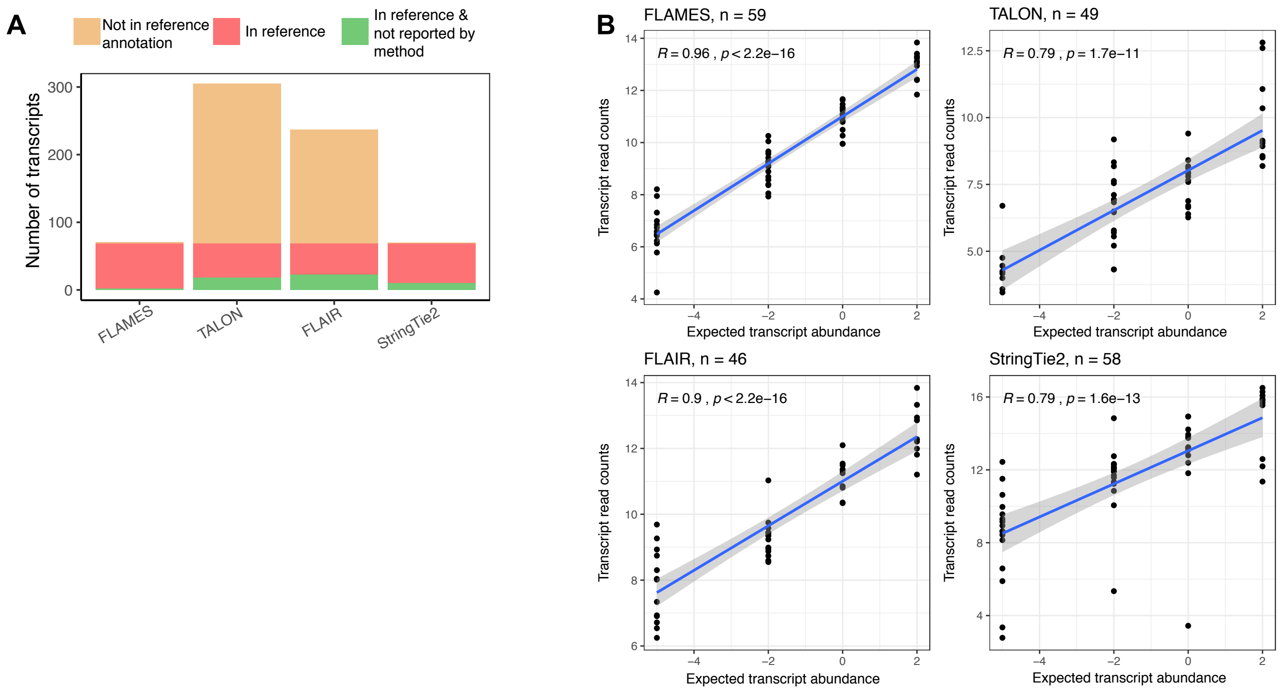


**Figure S2. Comparing *FLAMES* with other methods using the SIRV dataset.**
(A) Number of transcripts detected and not detected by each method, colored by whether transcript is present in the reference annotation.
(B) Scatter plot of transcript quantification where the x-axis is the expected transcript abundance based on the known concentration in the spike-in sample and the y-axis is the recovered transcript count ($\log_2$) generated by each method. The Spearman correlation coefficient (R) and *P*-value and the number of transcripts (*n*) included in the analysis are annotated in each plot.

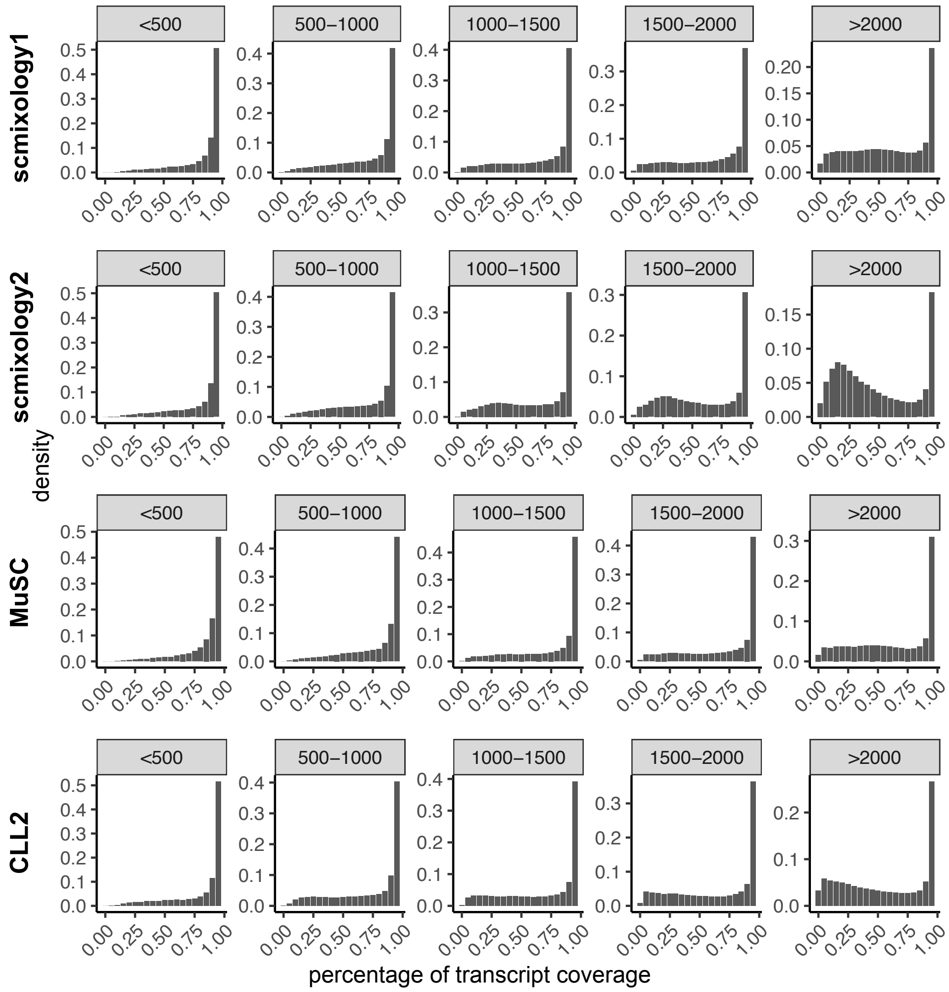


**Figure S3. Percentage of transcript coverage of individual reads stratified by length.**
The histogram shows the distribution of raw transcript coverage of reads aligned to the transcripts generated by *FLAMES*. Transcripts are stratified by length and divided into equal 500bp bins.

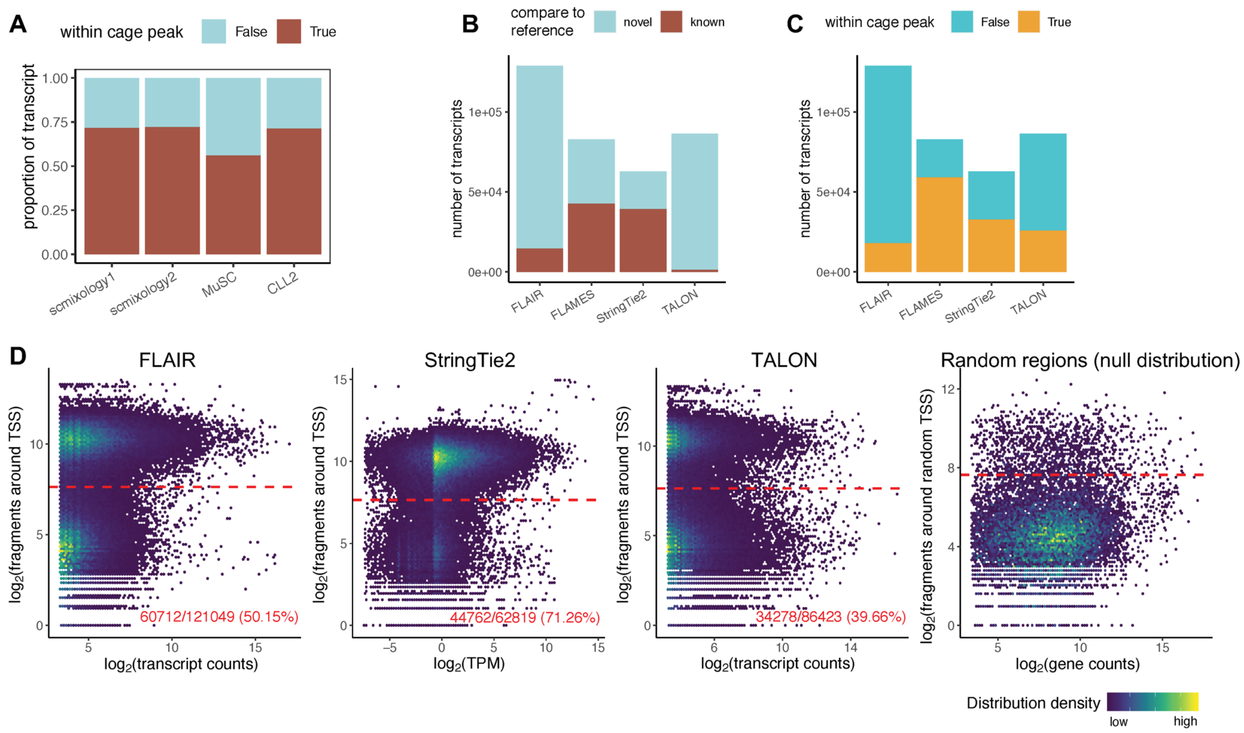


**Figure S4. Comparing *FLAMES* results with database annotations and other methods.**

(A) Proportion of transcript for which the TSS is within the FANTOM cage peak annotations for each sample.

(B) Number of transcripts generated by each method and the number of transcripts that match to the reference annotation.

(C) Similar to (B) and colored by the number of transcripts that fall within the FANTOM cage pack annotations.

(D) From left to right, density scatter plot showing total transcript count versus scATAC-seq read count around the TSS regions for *FLAIR*, *StringTie2* and *TALON*, together with a similar plot based on random TSS. The red line shows the threshold that separate the open chromatin and the background. The percentage shows the transcripts that have their TSSs in open chromatin regions.

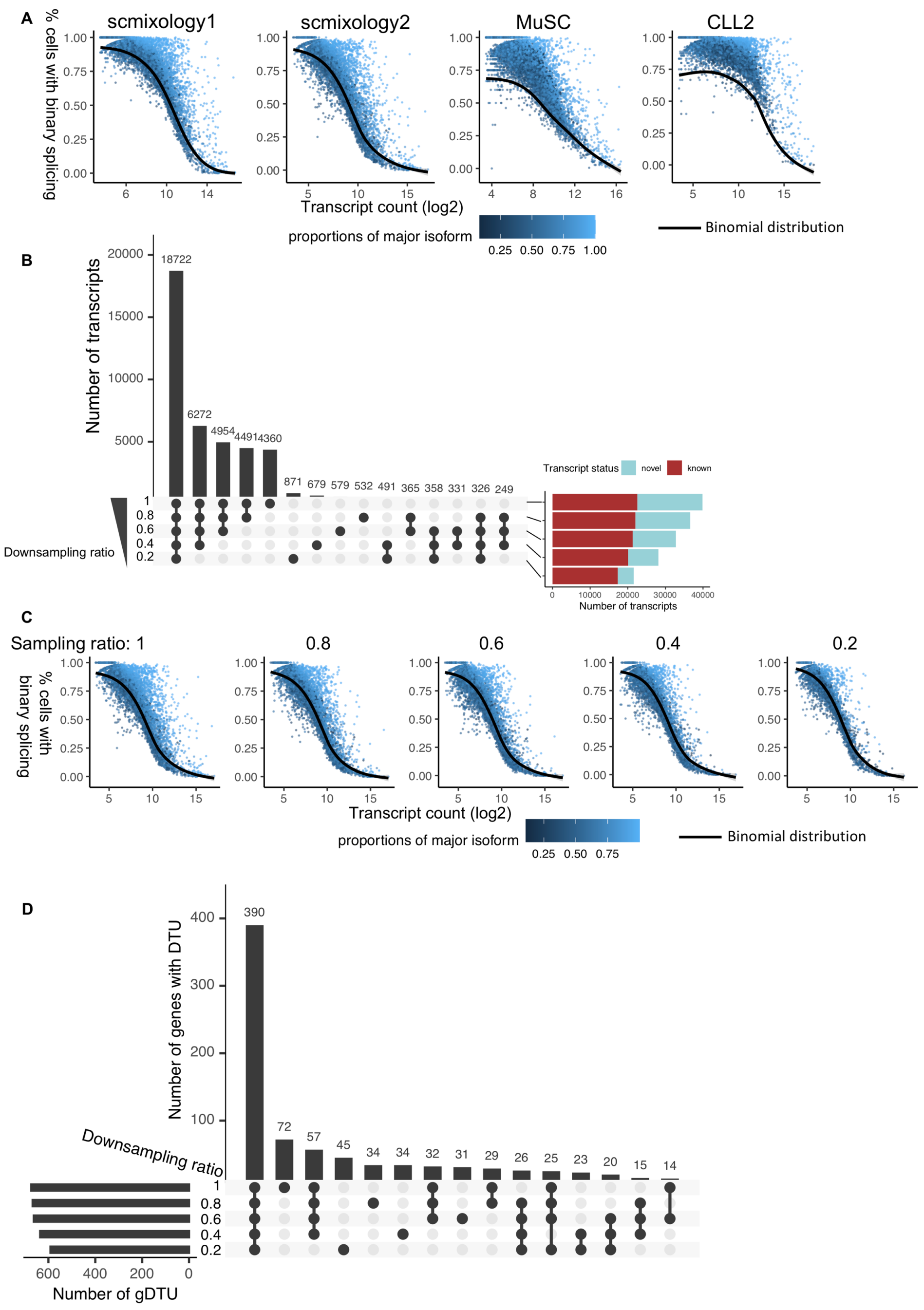


**Figure S5. Binary splicing and downsampling analysis.**
(A) scatter plot shows the total transcript counts for each gene versus the percentage of binary splicing, colored by the percentage of major isoform counts. Black line shows the null distribution that follows binomial distribution.
(B) Upset plot shows the overlap of transcripts generated from downsampling analysis of *scmixology2* with sampling ratio from 0.2 to 0.8, annotated with number of transcripts match to reference annotation.
(C) Similar to (A), scatter shows the binary splicing of samplings with different downsampling ratio of *scmixology2*.
(D) Upset plot shows the overlap of genes with significant DTU on *scmixology* data with different downsampling ratios.

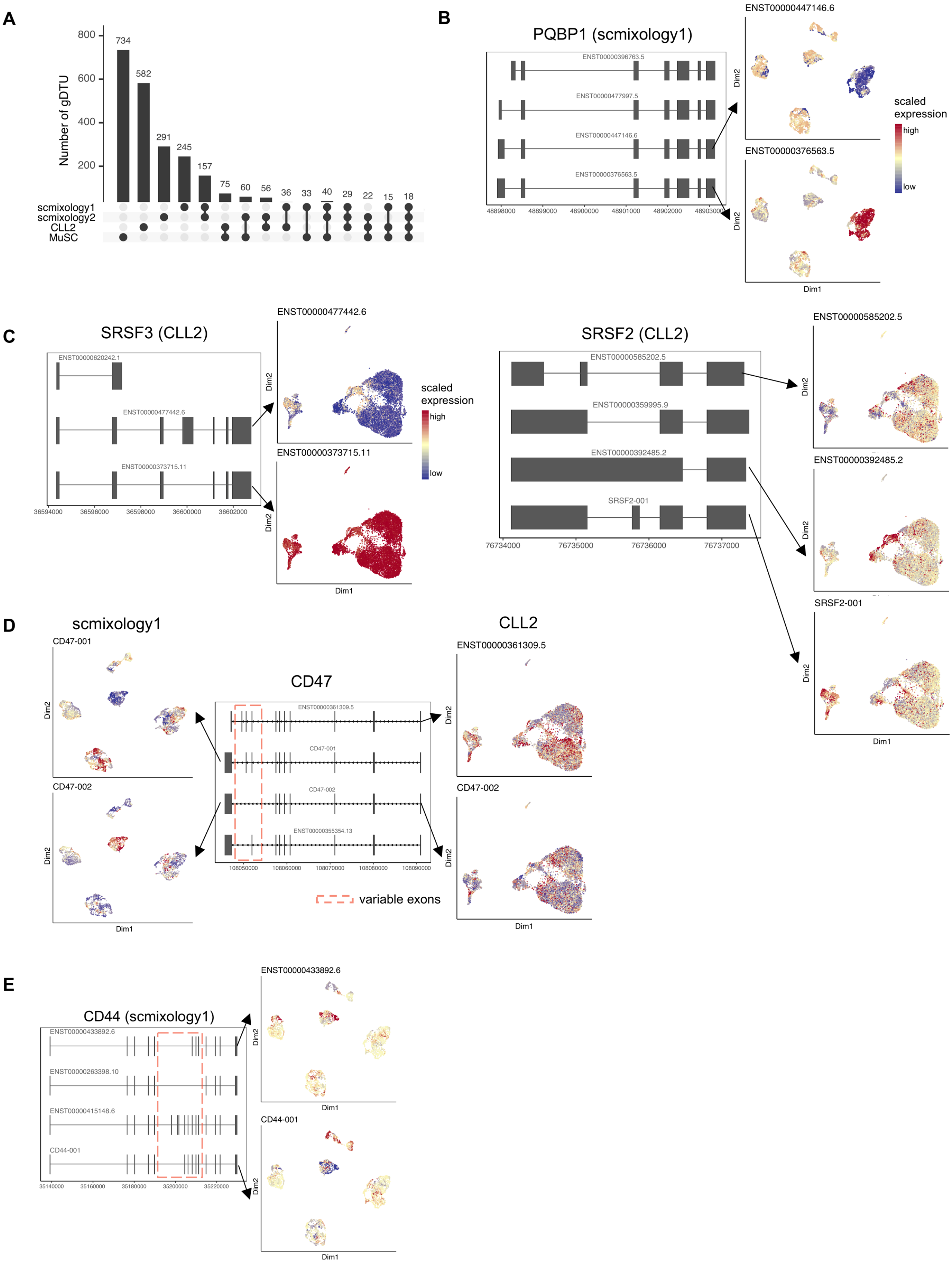


**Figure S6. Genes with differential transcript usage.**
(A) Upset plot shows the overlap of genes with differential transcript usage (gDTU) across different samples.

(B) Top 4 most abundant isoforms of PQBP1 in *scmixology1* and UMAP visualization colored by the expression of two isoforms with differential expression across different cell lines.
(C) Isoforms of SRSF3 in *CLL2*, with UMAP visualization colored by two isoforms with differential expression across different clusters
(D) Top 4 most abundant isoforms of CD47 in *scmixology1* and *CLL2*, together with UMAP visualization of expression of selected transcripts. Exons with alternative splicing was highlighted.
(E) Top 4 most abundant isoforms of CD44 in *scmixology1*, and UMAP visualization of transcript expression.

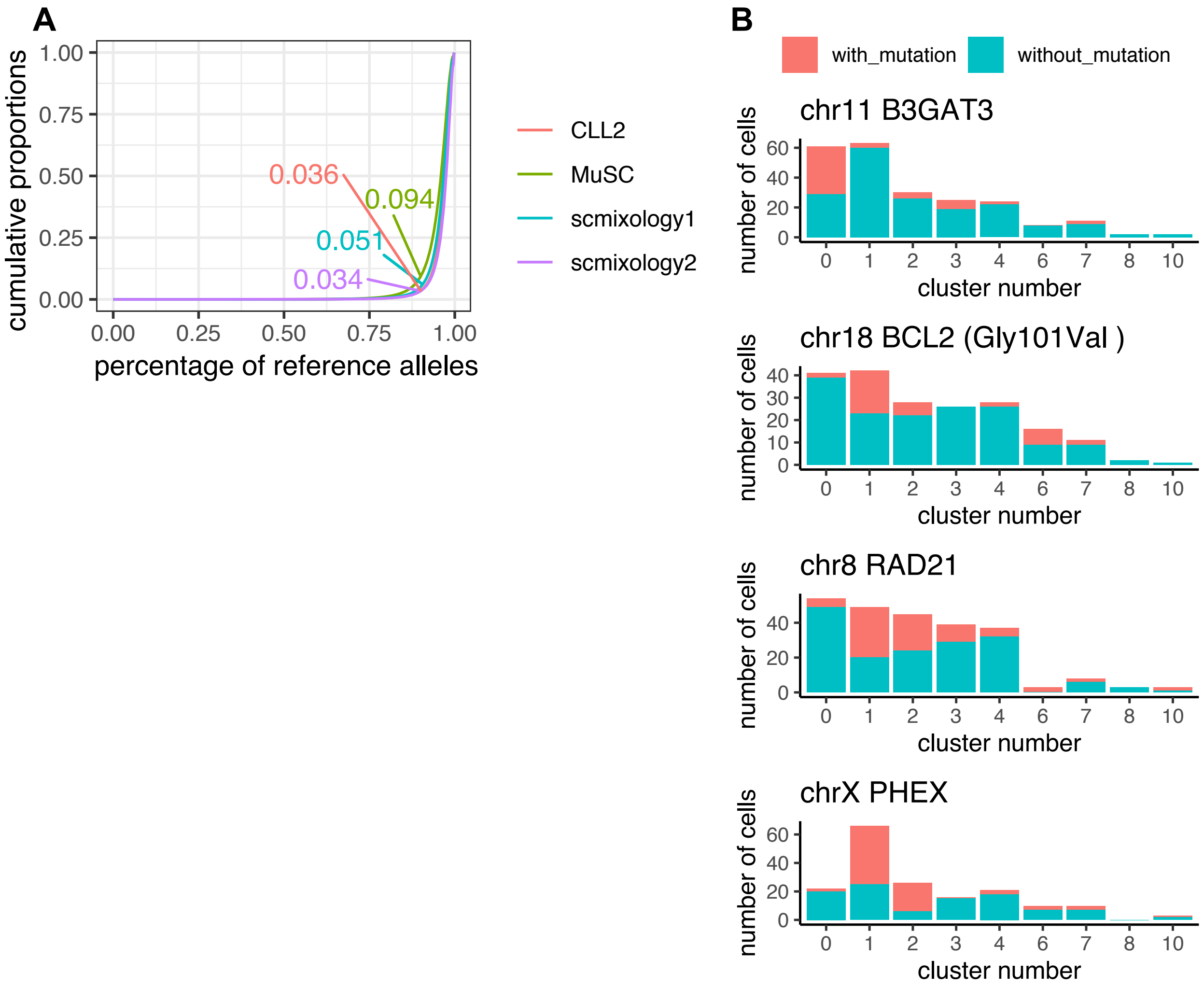


**Figure S7. Identification of genomic variation from long read by *FLAMES*.**
(A) Cumulative proportions of bases with certain percentage of reference alleles. The cumulative proportions of bases with 90% allele frequency were labelled on the plot, represent the percentage of reads that have lower than 90% reference allele frequency. (B) Bar plot of four significant mutations shows the distribution of cells in each cluster (x axis) that detected the mutation.
